# Touch DNA on objects can be analysed at low cost using simplified direct amplification methods

**DOI:** 10.1101/540823

**Authors:** Katherine Gammon, Kirk Murray-Jones, Daniel Shenton, Zoe Wood, Carl Mayers

**Affiliations:** Dstl, Porton Down, Salisbury, SP4 0JQ, UK

**Keywords:** touch DNA, direct amplification, forensic DNA analysis, trace evidence

## Abstract

Previous studies reported in the literature demonstrate that a range of sampling vehicles can be used effectively for forensic analysis of human DNA in direct amplification reactions. In this study we compared Copan microFLOQ^®^ swabs with a range of alternative sampling vehicles, using touch DNA samples donated by 15 different volunteers. MicroFLOQ swabs performed well, as did 3 mm diameter discs punched from analytical filter paper. The 3 mm discs could be used in a 5 µl PCR volume, increasing sensitivity, and reducing costs when compared with other methods that require a larger PCR volume. Other inert sampling vehicles, such as interdental toothbrushes and toothpicks also gave good results in direct amplification. The study found a large variation in results between the 15 touch DNA donors, demonstrating the importance of validating touch DNA recovery techniques with a large pool of donors.

## Introduction

Touch DNA is a term used to describe human DNA that is left behind on an object after it has been handled. Forensic DNA technology has now advanced to the point where a successful DNA profile can be generated from the DNA of a single human cell [1]. The ability to successfully analyse such minute quantities of DNA increases the chances of generating useful forensic information from touch DNA. Evidence produced from the analysis of trace amounts of DNA is admissible in court, from sampling of gloves for traces of shed skin [2] to bite marks for salivary DNA [3].

Touch DNA analysis techniques vary according to the type of substrate which needs to be analysed. Tape sampling is generally favoured for ridged or absorbent surfaces, while swab sampling is more suitable for smooth surfaces, non-adsorbent surfaces and heavily shedding cloth. Further improvement in DNA recovery from surfaces can be achieved using swab and solvent pairing, with non-polar surfactants being well suited for this role as they do not interfere with downstream DNA amplification [4].

The processes involved in forensic DNA analysis usually include DNA extraction, quantification and short tandem repeat (STR) amplification. The pre-amplification processes of extraction and quantification consume a proportion of the original sample. This reduces the amount available for analysis, potentially causing the loss of valuable evidence. While DNA extraction removes cellular debris and any PCR-inhibiting compounds co-deposited or introduced during sampling, it is an inefficient process, leading to loss of sample. Many extraction methods are available, which vary in the efficiency of their retention of the original DNA template, with reported losses between 20 and 90% of the original template [5]. The loss of template during the extraction and purification procedure has a profound impact on the success of generating useful profile information from trace DNA samples.

Direct PCR is used extensively in molecular biology as a technique to introduce samples directly into amplification reactions without any prior DNA extraction and purification. As touch DNA samples are typically very low biomass samples by nature, their introduction directly into STR amplification is desirable to avoid the sample loss inherent in extraction processes. A further benefit of this approach is to reduce the chance of sample contamination by reducing the amount of handling needed. These benefits combine to maximise the chances of obtaining usable DNA profiles from touch DNA.

Direct PCR of trace DNA can be performed by the direct introduction of the substrate into the PCR reaction, where the substrate is such that it can be cut into small enough pieces, i.e. fabric [6]. Alternatively, swabs or other materials can be used to remove touch DNA from a substrate and then the swab head can be introduced directly into the PCR reaction. The successful use of direct amplification for touch DNA recovery has been widely reported from a range of sample types [7-11].

We have historically found direct amplification to be a useful technique for amplifying touch DNA sampled from objects, and have had good results using both neonatal flocked nylon swabs, and small cellulose discs punched from analytical grade filter paper [12], [13]. Wetted neonatal flocked swabs [12] are very effective at removing touch DNA from objects, and are compatible with direct amplification reactions. However, a major disadvantage of neonatal swabs is their large size, requiring a 50 µl PCR volume in order to ensure the entire swab tip was entirely submerged during amplification.

Commercial products designed for direct amplification are now available, notably microFLOQ^®^ swabs from Copan Diagnostics. Reports of touch DNA sampling with microFLOQ swabs have been very promising [14]. We report here a trial on direct amplification of touch DNA taken from volunteers, carried out in order to evaluate microFLOQ swabs for touch DNA, and compare them with both proven and experimental direct amplification techniques.

## Materials and methods

### Reference and control samples

Buccal cell and touch DNA samples from individuals were collected and anonymised in accordance with the Human Tissue Act 2004. Buccal cell samples were collected by swabbing the inside of the cheek with a sterile cotton swab (Copan Diagnostics, USA) for 30 seconds followed by deposition onto a Whatman™ indicating FTA^®^ card (GE Healthcare Life Sciences, UK). A 1.2 mm punch from each FTA card was added to 15 µl of low TE buffer (10 mM Tris-HCL, 0.1 mM EDTA) in a 0.2 ml PCR tube. GlobalFiler™ PCR reagents were prepared as per manufacturer’s instructions (Thermo Fisher Scientific, USA) and 10 µl added to the tubes containing samples in order to give a final reaction volume of 25 µl. PCR amplification was carried out as per manufacturer’s instructions, but with the cycle number adjusted to 26 cycles. Capillary electrophoresis was carried out using a 3500xl Genetic Analyzer (Thermo Fisher Scientific) and data analysed using GeneMapper™ ID-X v1.4 (Thermo Fisher Scientific). SRM2391c control DNA was obtained from the National Institute of Standards and Technology (USA).

### Collection of touch DNA samples

Swatches of black polyurethane plastic (2 × 8 cm) were subjected to two rounds of ethylene oxide sterilisation in order to ensure they were free from any DNA contamination. Touch samples were collected from volunteers who had been instructed to wash their hands 30 minutes prior to sampling. Volunteers were asked to handle a swatch for 60 seconds, rubbing both sides of each swatch in turn. Swatches were individually bagged and stored at room temperature for a maximum of four weeks prior to DNA recovery. To recover touch DNA, both sides of each swatch was divided into eight 10 × 10 mm sampling regions. The sampling techniques described below were tested in groups, on independent swatches produced by different volunteers.

### Recovery of touch DNA using 4N6 flocked nylon swabs and EZ-1 extraction (control method)

In order to provide a control method, a sample from each tested swatch was collected using a 4N6 FLOQSwab^®^ (Copan Diagnostics) and DNA extracted using a Qiagen EZ-1 instrument (Qiagen, Netherlands). One side of a 4N6 FLOQSwab was wetted with 2 μl of molecular biology grade water and rubbed over the surface of the swatch, followed by the dry side of the swab for a total of 15 seconds. Swabs were dried at room temperature for 10 minutes, and DNA extracted using a Qiagen EZ-1 instrument using the DNA Investigator kit (Qiagen). Swab heads were processed according to Qiagen protocols, snapping the tip into a 2 ml tube containing 290 µl G2 buffer and 10 µl proteinase K and incubating at 56 °C for 15 minutes. Prepared swabs were then extracted in the EZ1 using the tip dance protocol, with DNA eluted into 40 µl water.

### Recovery of touch DNA using wetted microFLOQ® swabs

A microFLOQ Direct swab (Copan Diagnostics) was wetted with 1 μl of molecular biology grade water and used immediately to rub the sampling area for 15 seconds. The swab tip was then snapped off into a 0.2 ml tube containing PCR reagents diluted to working concentration. In addition to using molecular biology grade water, other wetting agents were also tested. These were 1 μl volumes of either 2 % (v/v) Triton X-100 (Sigma-Aldrich, USA), or 5 % (w/v) Brij^®^ 58 (Sigma-Aldrich).

### Recovery of touch DNA using microFLOQ swab wetted with SpeedBeads™

SpeedBeads magnetic carboxylate modified particles (Sigma-Aldrich) were washed by adding 1 ml of molecular grade water to 20 µl of beads and vortex mixing for 10 seconds. The beads were sedimented using a magnet, the supernatant removed, and beads re-suspended in 20 µl molecular grade water. The bead mixture was vortex mixed for 30 seconds immediately prior to use. For sampling, a microFLOQ® Direct swab was moistened with 1 μl of the SpeedBeads mix, then used immediately to rub the sampling area with the tip of the swab for 15 seconds. The swab tip was then snapped off into a 0.2 ml tube containing PCR mastermix.

### Recovery of touch DNA using cellulose discs

A paper punch was used to make 3 mm diameter circles of Whatman™ Grade 1 180 μm thick cellulose filter paper (GE Healthcare Life Sciences), which were stored in sterile disposable petri dishes. Using a sterile 20 µl pipette tip a disc was picked up and placed onto the sampling area and 2 µl of molecular grade water was pipetted directly onto the disc. Using a fresh pipette tip the disc was rubbed around the sampling area for a total of 15 seconds and then added directly to a 0.2 ml tube containing PCR mastermix. Examples of sampling methods used are shown in Figure 1.

**Figure 1.**
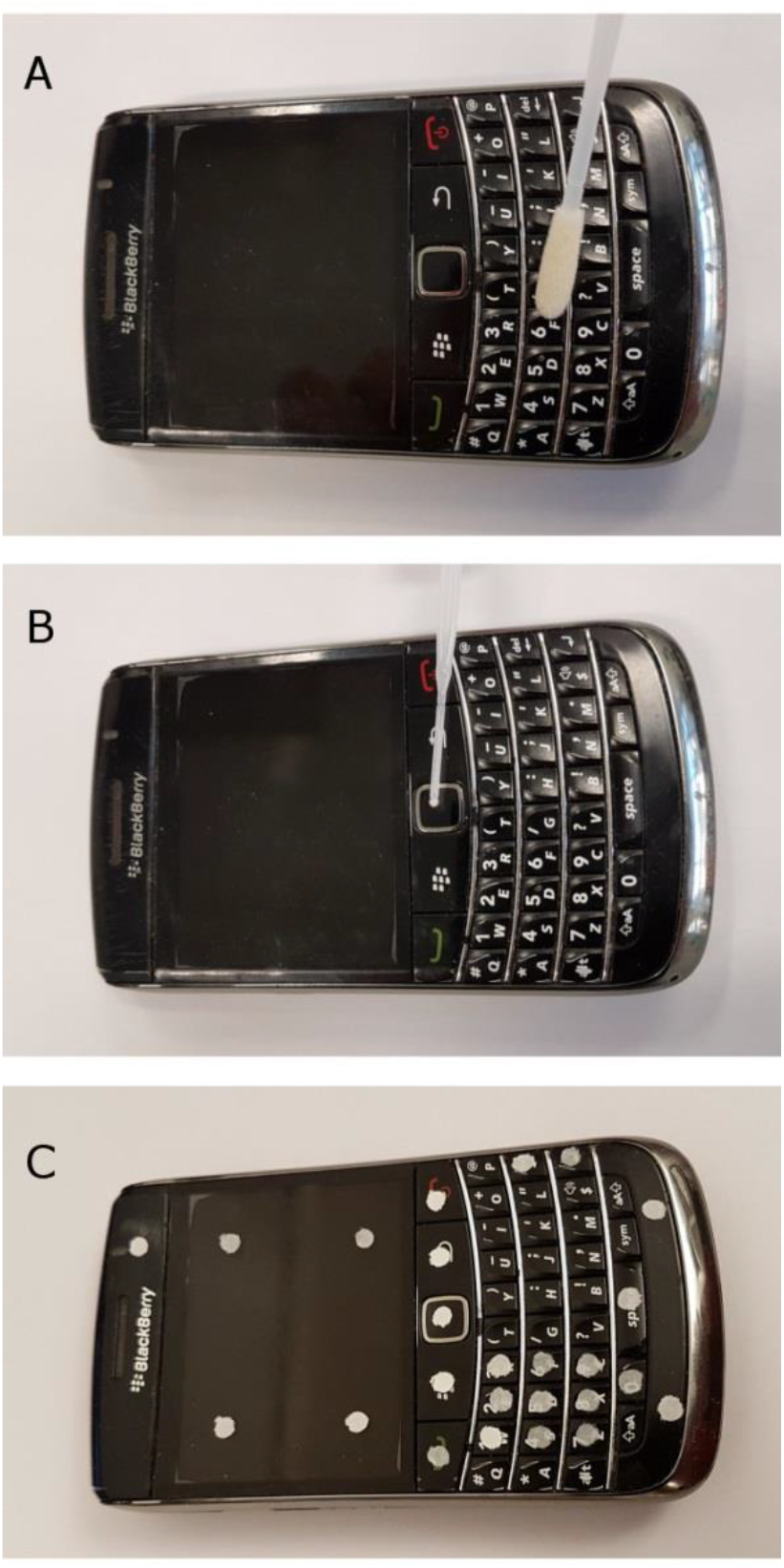
Examples of sampling methods used in this study. Touch DNA samples were sampled (illustrated here with a BlackBerry™ mobile phone for scale) using a) Copan 4N6 FLOQ™ swab, b) Copan microFLOQ^®^ swab, and c) 3 mm discs of Whatman filter paper.

### Recovery of touch DNA by elution in liquids

In order to test the recovery of touch DNA into an aliquot of water, 10 µl of molecular grade water was pipetted directly onto the sample area and left undisturbed for 30 seconds. The water was then aspirated by pipetting up and down 5 times using a pipettor set to 7.5 µl. The final aspiration of 7.5 µl was added directly to a tube containing PCR mastermix. Other liquids were also tested in place of water. These were 2% (v/v) Triton X-100, 5% (w/v) Brij 58, and prepared GlobalFiler PCR mastermix.

### Recovery of touch DNA using interdental brushes

Interdental brushes, 0.4 mm diameter (TePe, Sweden), were tested as an alternative to microFLOQ swabs. A 2 µl volume of molecular grade water was added to the centre of the sampling area and the side of a 0.4 mm wire size interdental brush was used to scrub the sampling area for 15 seconds at a 45 degree angle. The head of the brush was cut into a 0.2 ml tube containing PCR mastermix using sharp scissors. Scissors were cleaned prior to and following use by submersion in Virkon^®^ solution (DuPont, USA) for 15 minutes followed by 30 minutes exposure in a UVP CL-1000 UV cross linker (Analytik Jena, USA).

### Recovery of touch DNA using silicone toothpicks

An EasyPick silicone toothpick (TePe, Sweden) was used to scrape over the sampling area for 15 seconds using both sides of the bottom half of the pick. The bottom 5 mm of the toothpick was subsequently cut into a 0.2 ml tube containing PCR mastermix using sharp scissors. Scissors were cleaned prior and following use as previously described.

### PCR amplification of touch DNA samples

In the case of the control EZ-1 DNA extract, 15 µl of extract was added directly to 10 µl of GlobalFiler™ PCR mastermix prior to amplification. In the case of direct amplification reactions, reactions were amplified in volumes of PCR mastermix ranging between 5 - 50 µl. All PCR reactions were amplified on an Applied Biosystems Veriti system (Thermo Fisher) for 29 cycles (Identifiler Direct) or 30 cycles (GlobalFiler), following manufacturer’s instructions.

### Capillary electrophoresis and analysis

Amplified products were processed on a 3500xL Genetic Analyzer, using 1 μl PCR product in 9.6 μl Hi-Di™ formamide containing 0.4 μl GeneScan™ 600 LIZ^®^ Size Standard v2.0 (Thermo Fisher Scientific). One microliter of allelic ladder was included per 24 sample injection. Plates were denatured at 95 °C for 3 minutes prior to loading on the 3500xL instrument. Electrophoresis was performed on a 36-cm capillary array with POP-4™ polymer (Thermo Fisher Scientific) using an injection time of 24 seconds. STR peaks were sized and typed using GeneMapper^®^ ID-X Software Version 1.4 (Thermo Fisher Scientific) using internally verified analytical thresholds. Alleles called by GeneMapper were compared with the reference profile for the donor. The number of alleles matching the reference profile was recorded, and divided by the number of peaks present in the reference profile, to give a percentage profile match of the experimentally-sampled touch DNA against the full donor profile from a buccal sample.

## Results and Discussion

The cellulose disc direct amplification method had previously shown greater sensitivity than the neonatal swab direct amplification method [12]. We speculated that this was due to the reduced PCR volume of the cellulose disc method (10 ul) compared with the neonatal swab method (50 ul). A five-fold reduction in PCR volume would cause a five-fold increase in DNA concentration in the amplification reaction, allowing for more efficient amplification with low copy number touch samples. When 100 pg of NIST SRM2391c reference DNA was dried onto replicate 3 mm diameter cellulose discs and subjected to direct amplification it was clear that a reduction in PCR volume gives much greater sensitivity (Figure 2). A 50 ul reaction containing a disc with 100 pg of reference DNA only produced a median of 4% of profile recovery (n=9), while the same 100 pg of reference DNA in a 5 ul reaction produced a median of 45% profile recovery (n=9).

**Figure 2.**
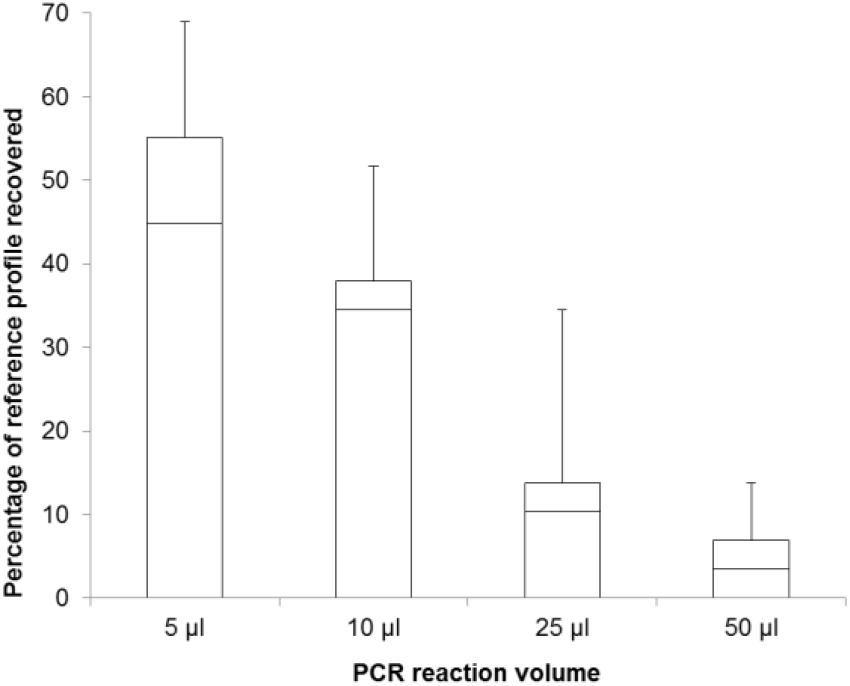
Percentage of reference profile recovered from a 3 mm cellulose disc seeded with 100 pg DNA, amplified in different PCR reaction volumes. The efficiency of amplification in different PCR reaction volumes is shown as the percentage of reference profile alleles recovered in a direct-amplification reaction using the Applied Biosystems IdentiFiler™ multiplex. Each PCR reaction included a 3 mm cellulose disc seeded with 100 pg of NIST SRM2391c reference DNA. Results are shown as box-and-whisker plots; median is marked as a horizontal line, boxes indicate 25th to 75th percentile, whiskers indicate minimum and maximum values (n=9 amplification reactions).

These results suggested that other types of swabbing methods and sampling vehicles could also work well in direct amplification reactions. This was investigated using plastic swatches that had been handled by volunteers (Figure 3). As a control, Copan 4N6 flocked nylon swabs were used to sample the plastic swatches, and DNA was extracted using a Qiagen EZ-1 instrument. Attempts were also made to pipette liquid directly onto the surface, recovering the liquid with a pipette tip in an attempt to wash trace DNA off the surface. None of these liquid methods worked well (Figure 3), despite adding surfactants [2% (v/v) Triton X-100 or 5% (v/v) Brij]. Pipetting PCR master mix onto the surface, followed by direct amplification also proved to be ineffective compared with the standard EZ-1 extraction method.

**Figure 3.**
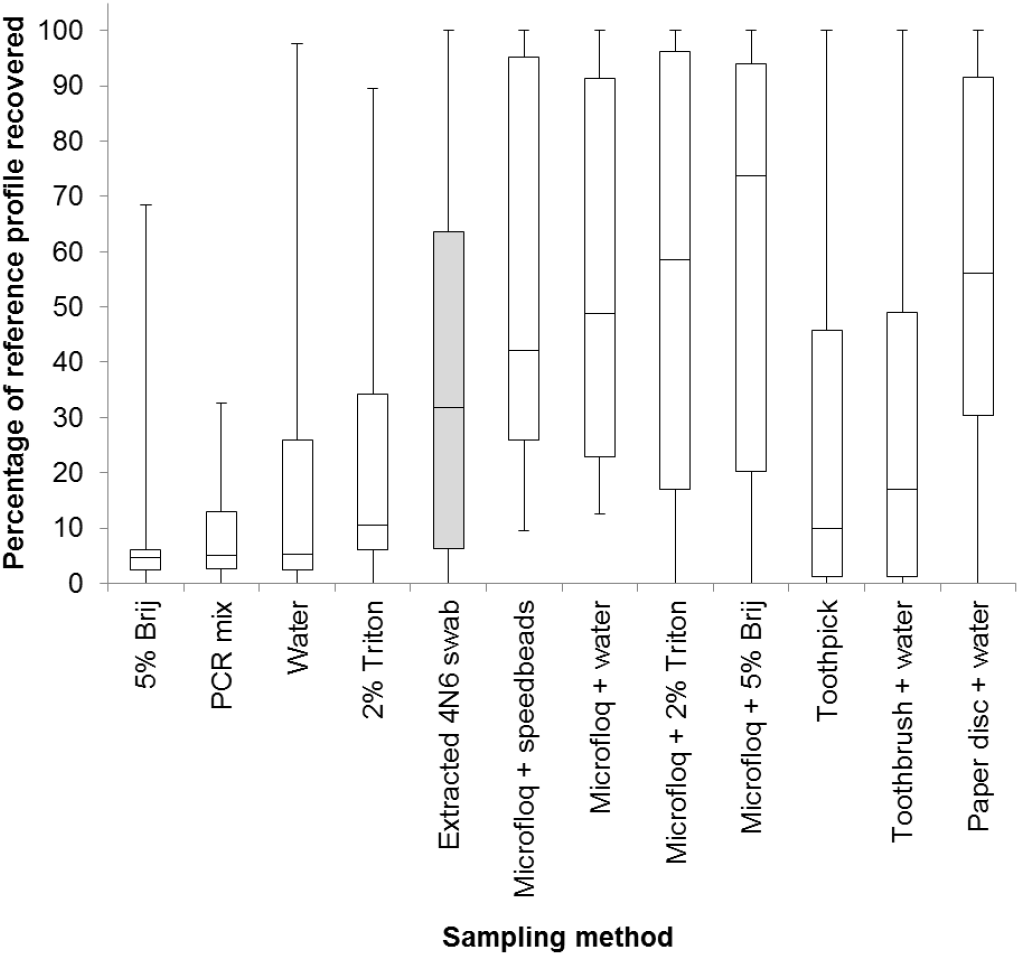
Percentage of reference profile recovered using different direct amplification methods. The efficiency of touch DNA profile recovery is shown as the percentage of donor reference alleles recovered in a single 12.5 ul direct-amplification reaction using the Applied Biosystems GlobalFiler multiplex. Each direct amplification method was tested on touch samples provided by 15 different touch DNA donors. Results are shown as box-and-whisker plots; median is marked as a horizontal line, boxes indicate 25th to 75th percentile, whiskers indicate minimum and maximum values (n=15 volunteer samples per method). Shaded box indicates the control method (Copan 4N6 FLOQ swab extracted using Qiagen EZ-1).

MicroFLOQ swabs and 3 mm cellulose discs both performed well in a 12.5 ul direct amplification reaction, giving results comparable to the control. Addition of surfactants [2% (v/v) Triton X-100 or 5% (v/v) Brij] was tested to see if they would increase the yield of touch DNA removed from the surface, but did not greatly change the sensitivity. SpeedBeads were added to the swabs to test if this would enhance the removal of material from the surface due to their scouring action, but also did not greatly change the sensitivity. Rubberised toothpicks and interdental toothbrushes can prove useful in sampling crevices on objects, and both gave usable results in direct amplification reactions (Figure 3).

There was a large variation in the results from the 15 donors used in this study. In order to examine the repeatability of microFLOQ swabs in direct amplification, four repeat microFLOQ swabs were used to sample different areas of the swatches touched by each volunteer. While there was some variation within each group of four samples from each volunteer, there was very large variation between volunteers (Figure 4). Swatches handled by volunteer ‘O’ never generated more than a 25% profile, while swatches handled by volunteer ‘F’ always gave a 100% profile.

**Figure 4.**
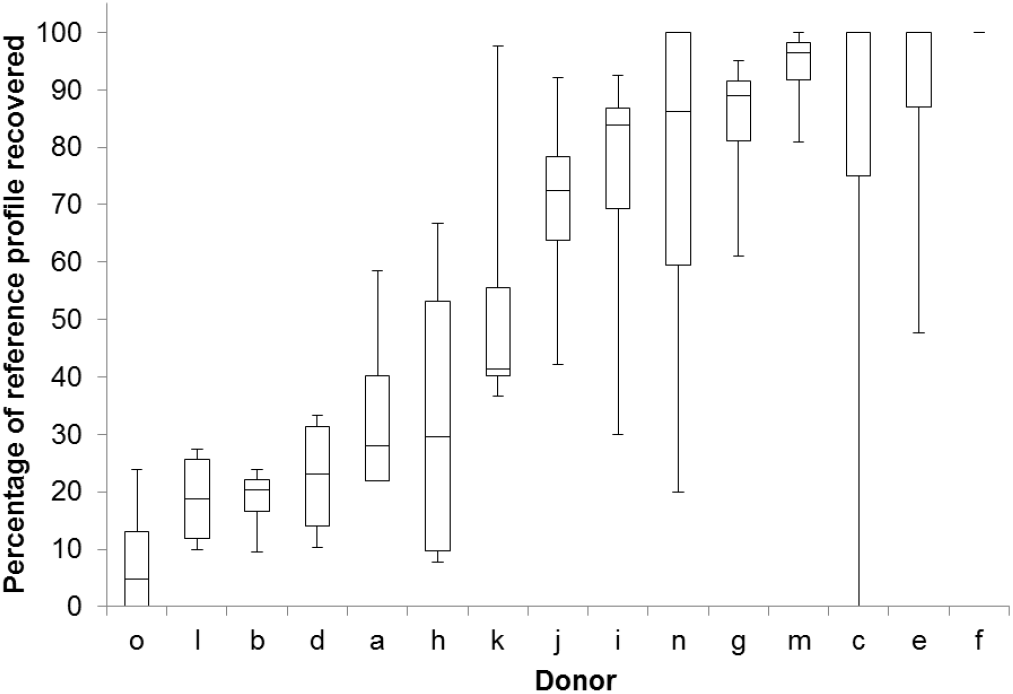
Percentage of reference profile recovered from touch samples supplied by different donors, using MicroFLOQ swabs in direct amplification. The efficiency of touch DNA profile recovery is shown as the percentage of donor reference alleles recovered in a single direct-amplification reaction using the Applied Biosystems GlobalFiler™ multiplex. MicroFLOQ direct amplification method was tested on touch samples provided by 15 different touch DNA donors. Results are shown as box-and-whisker plots; median is marked as a horizontal bar, boxes indicate 25th to 75th percentile, whiskers indicate minimum and maximum values (n=4 repeats per donor).

Direct amplification proved to be an economical and effective way to amplify trace DNA profiles from touched items. The process of direct amplification is by its nature more efficient; DNA cannot be lost from the sample as the entire sample is transferred to the PCR amplification reaction. Costs are greatly reduced due to the elimination of the extraction and quantification stages. Furthermore, use of a small sampling vehicle with a low volume allows a reduction in volume of the PCR amplification reaction. The use of 3 mm cellulose discs was particularly effective in a 5 ul PCR volume, cutting reagent costs significantly compared with the commonly used 12.5 or 25 ul volumes. The use of small sampling vehicles such as MicroFLOQ swabs or cellulose discs allows very small areas to be sampled discretely and precisely, reducing the incidence of mixtures from multiple donors.

Cellulose discs can be produced at very low cost, and represent an efficient way to sample touch DNA that is compatible with very small volume amplification reactions. In our experience, cellulose discs are best suited for laboratory sampling, while MicroFLOQ swabs are better suited to field situations. We have found that many inert sampling vehicles can be used in direct amplification reactions, as demonstrated in the examples shown here (rubberised toothpicks and interdental toothbrushes). We would like to encourage further experimentation with direct amplification; the method can be extremely sensitive, and significantly reduces costs and the incidence of mixtures. We suggest researchers should verify new methods using a large panel of volunteers as a source of touch DNA. Even with a proven sampling vehicle such as microFLOQ swabs we have found extremely variable results (Figure 4) between volunteer samples, and testing small numbers of samples can easily produce very misleading results.

## Acknowledgements

We would like to thank Penny Brookes, Sue Carolin, Nick Luck and Martin Pearce for valuable advice, review and support. This work was funded by the UK Ministry of Defence. The authors declare that no competing interests exist. © Crown Copyright (2019), Dstl.

